# Weed regulation by crop and grassland competition: critical biomass level and persistence rate

**DOI:** 10.1101/572701

**Authors:** Mauricio Z. Schuster, François Gastal, Diana Doisy, Xavier Charrier, Anibal de Moraes, Safia Médiène, Corentin M. Barbu

**Affiliations:** Department of Crop Production and Protection, Federal University of Paraná, 1540 Rua dos Funcionários Road, Curitiba, PR 80035-060, Brazil; INRA (Institut National de la Recherche Agronomique) UE1373, FERLUS, 86600 Lusignan, France; Agronomy, INRA, AgroParisTech, Université Paris-Saclay, 78850 Thiverval-Grignon, France

**Keywords:** weed management, weed ecology, weed biocontrol, weed modelling, weed population dynamics

## Abstract

1. It is widely agreed that competition regulates plant populations and shapes communities. Many studies have suggested that crop and grassland competition can be used for cost-effective sustainable weed control. However, effective weed management requires a precise knowledge of the effects of agronomic practices and there is a lack of quantitative indicators to compare and predict the success of weed biocontrol by competition.
2. We studied weed abundance dynamics over a 12-year period in crop-grassland rotations (rotation treatments consisted of maize, wheat and barley crops, alternating with temporary grassland maintained for three or six years in the rotation and fertilised with two different levels of nitrogen). In addition to classical statistical analysis of the different aforementioned rotation treatments, we also modelled weed abundance as a function of the crop and grassland competition, expressed here by biomasses harvested in the preceding years.
3. We show that weed abundance decreases over the years in grassland and subsequent crops only if the grassland receives sufficient nitrogen fertiliser. Our model had a much greater explanatory power than the rotation treatments. This model estimates a critical biomass level above which weeds are suppressed in subsequent years, and below which they tend to thrive. This critical biomass level was 24.3 and 4.7 tonnes ha^−1^ of dry matter for crops and grassland, respectively, highlighting the greater competitiveness of grasslands than of crops. Several clear differences between weed functional groups emerged.
4. *Synthesis and applications* - This new modelling approach directly links the interannual dynamics of weed populations to current and previous biomass production levels. This approach facilitates the development of environment-friendly weed management strategies and paves the way for comparisons of the competitiveness against weeds of crops and grassland under various pedoclimatic conditions and agronomic practices.

## 1. Introduction

Most food and feed production systems worldwide make use of synthetic herbicides for weed management. In this context, herbicide use has resulted in serious environmental and ecological problems (Boutin et al., 2014). Highly effective environment-friendly alternatives to chemical weed control, such as the use of crop and grassland competition with weeds, could potentially reconcile agricultural production and environment quality and play a key role in ensure global food security in the future (Petit et al., 2018; Gaba et al., 2018).

Many previous studies have shown how the manipulation of agronomic practices (e.g. seed rate, crop cultivar and row spacing and direction) to improve the competitiveness of the crop can help to control weeds (Sardana et al., 2017). Other studies have suggested that grassland is more efficient than crops for weed suppression (Meiss et al., 2010b; Schuster et al., 2018). However, several studies have shown that, in dry conditions (Miller et al., 2015) or at high grazing intensities (Schuster et al., 2016), the introduction of grassland into the rotation can have deleterious effects on weed control. Assessments of the competitiveness of crops and grassland against weeds would help to explain these divergent results, but quantitative indicators for predicting the success of weed biocontrol and comparing competitiveness between studies are lacking. Furthermore, such studies are often based on a snapshot characterisation of the effects of crop or grassland competition on weeds, with very few considering the impact over multiple years.

It is difficult to gain a comprehensive understanding of the dynamics of crop and grassland competition against weeds, not only due to the interactions between the grassland or crop and all the intrinsic components of the weed species (e.g. life cycle, leaf and root type, growth habit; Gaba et al., 2014), but also due to the interactions between weeds, environment (e.g. time of emergence, growth rate, seed production; Cirillo et al., 2018) and the management (i.e. farmer’s decisions) to which fields are subjected (e.g. tillage regime, fertilisation rates, crop rotation; Colbach et al., 2014).

We studied weed abundance dynamics in crop-grassland rotations over a 12-year period, to determine whether and how weed abundance during the crop and grassland phases of the rotation changes with the duration and fertilisation of grasslands. We also developed a statistical model with an explicit translation of cultivated plant competitiveness against weeds. We adjusted this model according to weed abundance data and tested the hypothesis that the effects of the duration and fertilisation of grassland, and of rotation schemes and weather conditions over the years can be measured as variations in grassland and crop biomass production, used as an indicator of competitive potential. Finally, we characterised the effects of weed traits on the competitiveness of crops and grassland.

## 2. Materials and Methods

### 2.1 Description of the site, management and rotation treatments

The long-term study of cropping systems including temporary grasslands analysed here is part of SOERE ACBB (Observatory and Experimental System for Environmental Research - Agroecosystems, Biogeochemical Cycles, and Biodiversity) and is located at INRA, Lusignan, France (46°25’13” N; 0°07’29” E, 151 m above sea level). This site has an oceanic climate with a summer drought, a mean air temperature of 12°C and a mean annual precipitation of 750 mm. The soil is a rubefied brown earth on clay, with traces of ferruginous shell.

This experiment was started in 2005 and conducted over 12 years. The treatments studied are rotations of maize, wheat and barley alternating with grassland. The grasslands were sown in mid-September with a mixture of three grass species: perennial ryegrass (*Lolium perenne* cv. Milca: 5 kg.ha^−1^), tall fescue (*Festuca arundinacea* cv. Soni: 10 kg.ha^−1^) and orchard grass (*Dactylis glomerata* cv. Ludac: 12 kg.ha^−1^). Grasslands were mowed to a height of 5 to 7 cm and harvested three to five times per year, depending on climate and biomass production, and the cut grass was removed from the field. The first cut took place in the spring (April). Grasslands were fertilised after each cut (see Kunrath et al. 2015 for more details). The weed species present was described in Weed_traits_SuppInfo and their management have been described elsewhere (Doisy 2015). Briefly here, in order to control weed invasion, post-emergence herbicide was applied on annual crops every year during April. No herbicide was applied on grasslands, except once during grassland installation in 2005.

The five rotation treatments were distributed in four blocks of individual plots of 4000 m^2^ each (for more details see http://www.soere-acbb.com/demarche-experimentale). The rotation treatments consisted of: cereal-based rotation with repeated maize/wheat/barley sequences (C); a crop-grassland rotation, in which the cereal sequence was followed by three years of grassland (G3C); a grassland-cropping rotation, beginning with six years of grassland receiving high (∼230 kg ha^−1^ year) or low (∼30 kg ha^−1^ year) levels of nitrogen fertiliser (G6C and -G6C, respectively) followed by the cereal sequence; and a continuous grassland with high levels of nitrogen fertilization, as defined above, (G).

### 2.2 Data collection

#### 2.2.1 Field sampling to estimate weed abundance

In each experimental unit, from 2005 to 2017, weed abundance was determined in the cereal fields in April, before post-emergence herbicide application, and then again in early autumn, before crop harvest. In the grasslands, weed abundance was determined before the first cut (April). In each plot, 13 points were sampled at 12 m intervals along two 72 m transects laid out in an “X” pattern starting 5 m from the edge of the field. At each point, the abundance of each weed species was estimated with the Barralis scale adapted for an area of 0.25 m^2^, with classes “0” to “4” corresponding to, one, two to five, six to 12 and more than 12 individual weeds, respectively. When weed abundances were measured twice a year, we used the maximum abundance by species observed for each measurement point on both dates. Hereafter, the density per point for a group of weed species is the sum of the lowest abundance values of the Barralis interval for the species present at each point, expressed per m^2^.

#### 2.2.2 Field sampling to estimate crop and grassland biomass

Biomass production in the fields was estimated by harvesting with an experimental harvester equipped with an on-board weighing system (Haldrup, Germany), and drying to constant weight in an oven at 70°C to determine dry matter content per unit area (DM t ha^−1^). We estimated biomass production just before harvest, for both grassland and crops, at three different sampling points per field. Each sample covered an area of at least 7.50 m^2^, corresponding to the passage of a harvester with a 1.5 m cutting bar over a distance of 5 m. The length of the sampling area was increased if smaller amounts of biomass were produced or if biomass production was heterogeneous. The cutting bar was set at a height of 5 to 7 cm (the same height as for the mowing of grassland plots).

#### 2.2.3 Weather data

Yearly rainfall, mean air temperatures and thermal amplitude data at a height of 2 m were measured at the experimental site and are available from the Climatik database maintained by INRA AgroClim.

### 2.3 Data management and general modelling procedures

The abundance of weeds in a field plot was summarised as the sum of the abundances, expressed per m^2^, at the 13 sampling points. We modelled this total abundance by a negative binomial distribution with a log link (Bates et al., 2015). We compared the models with Akaike’s information criterion (AIC) in R regression packages (Sakamoto and Akaike, 1978). We checked that the residuals were not autocorrelated over time, using acf in the R package itsadug (van Rij et al. 2017). The significance of differences between factor levels, such as the different rotations and crops in place, according to the models obtained, was assessed in pairwise comparisons with an alpha risk of 0.05, with Holm–Bonferroni adjustment, as implemented in the R package lsmeans (Lenth, 2015). The relevance of the models for describing the variability in the data was assessed with Fisher’s “goodness of fit” test (Fisher, 1924).

### 2.4 Basal model of weed control by crop and grassland biomass

We propose a model connecting weed abundance to the biomass harvested in previous years and show how this model can be transformed into a generalised linear model that is easy to fit with statistical software. We assume that, for each year, if the harvested biomass *B* (i.e. the total of the three to five harvests over the year for grassland, and above-ground biomass at harvest for the crops; i.e. grains plus leaves and stems) is above a crop-specific critical level, *S*_*c*_, then weed abundance, *W*, tends to decrease the following year. By contrast, the weed flora was assumed to increase if the cultivated biomass (crop and grassland) was below *S*_*c*_. We account for this effect by considering the estimated abundance of weeds emerging in year T, 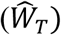, to be proportional to a power of the *S*_*c*_*/B* ratios of the preceding years.

We also considered ratios to have a decreasing impact on weed density over time (years), and we accounted for this decrease by modulating the ratios by a power coefficient inversely proportional to the number of years elapsed:

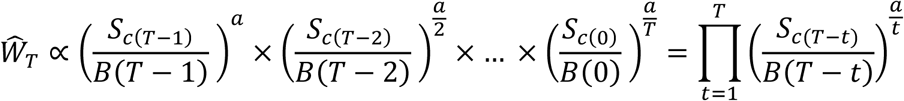

This can be expressed logarithmically, to obtain a linear formulation:

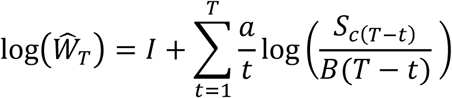

where I is an intercept corresponding to the logarithm of a basal level of weeds. In the above equation, the biomass over time and the inverse of time are separable:

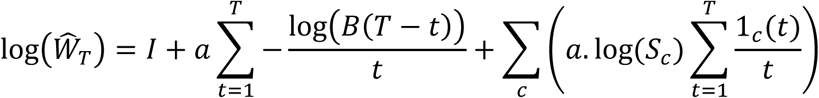

where 1_*c*_(*t*) is the indicator function for the presence of crop *c* in year t. The terms of this linear expression are identifiable with the terms of a negative binomial regression with a logarithmic link:

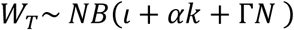

- where *ι* is the intercept of the regression corresponding to *I* in the initial linear expression. We use this intercept to account for the block of plots, *P*, and the current crop in the plot, *C*, which we subsequently treat as random effects.

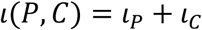
- *k* accounts for previously harvested biomasses: 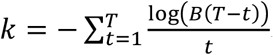, the sum of the log of the harvested biomasses inversely weighted by the time elapsed, multiplied by *α* = *a*, the corresponding coefficient in the regression.
- *N* is a vector of crop factors corresponding to the sum of the inverse of the time elapsed since the presence of the crop: 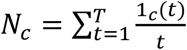, multiplied by Γ, the vector of the corresponding regression coefficients for each crop, with Γ_*c*_ = *a*.*log* (*S*_*c*_).

Once *k* and *N* have been calculated, the model can be fitted, with, for example, the glm.nb function of the R package MASS. The critical biomass level for each crop *c* is the exponential of the ratio of the regression coefficients:

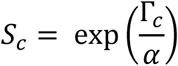

As the order of the cash crops in the rotation was always the same, it was not possible to determine critical biomass levels for individual crops. We simply distinguished grassland from cash crops, grouping maize, wheat and barley together to obtain a common *S*_*c*_ for these crops. The time required to reduce the impact of a given year under a given threshold is directly proportional to the coefficient α in the regression (see supplementary materials “alpha_visualization_SuppInfo.docx”).

We report the confidence intervals for *Sc* according to the exponentiation of Fieller’s confidence interval for a ratio of parameters (Fieller, 1954). The significance of the difference between parameter values estimated with different models was assessed by a million draws from the estimated normal distribution (for α) or transformed multinormal distributions of Γ_*c*_ and *α* (for *S*_*c*_), with comparison of the draws obtained, in pairs, for two different models or crops. If one of the parameters was greater than the other 95% of the time, we considered the parameters significantly different.

### 2.5 Variants of the basal model

We ran the model on subgroups of weeds defined on the basis of their common traits. Weeds were successively split into groups (see Weed_traits_SuppInfo) according to their life cycle (annual vs. perennial), taxonomic group (monocots vs. dicots), root structure (fibrous roots vs. tap roots), and growth habit (upright/climbing/rosette/creeping).

We used several indicators to characterise the quality of the predictions from the various models fitted: the root mean square error of prediction (RMSE) and bias, as implemented in the hydroGOF package (Zambrano-Bigiarini, 2014), and Pseudo-R^2^ and Spearman’s rank correlation as implemented in the Hmisc package (Frank and Harrell, 2016). Data assembly, consolidation and analysis were performed in R software for statistical computing version 3.1.3 (R Development Core Team, 2015). Detailed statistical codes are provided (CBL_PaperScript_R_SuppInfo and CBL_functions_R_SuppInfo).

## 3. Results

### 3.1 Effects of grassland duration and fertilisation on weed abundance

Weed abundance differed considerably between the crop-grassland rotations studied, and seemed to follow different trajectories over the years according to the management system (Figure 1A). Over the last 10 years of the study, mean weed abundance was systematically lower in the continuous grassland than in the cereal-based rotation (rotation treatments G vs. C). Mean weed abundance was generally lower in the rotation treatment including six years of well-fertilised temporary grassland than in the cereal-based rotation (Figure 1B, rotation treatments G6C vs. C). By contrast, six years of temporary grassland with low levels of nitrogen fertilisation in the rotation resulted in higher weed abundances, similar to those obtained in the rotation without grasslands (Figure 1B, rotation treatments -G6C and C, respectively). Rotation treatments including well-fertilised grassland maintained over a period of six years had lower weed abundance than rotation treatments including well-fertilised grassland maintained over only three years (rotation treatments G6C vs. CG3). Weed abundance declined progressively over successive years of the grassland phase in rotation treatments receiving high levels of nitrogen fertiliser (rotation treatments CG3, G6C and G, years 2005-10) but not in the rotation treatment in which the grassland received low levels of nitrogen fertiliser (rotation –G6C) (Figure 1A). We assessed the statistical significance of the differences in weed abundance between these rotation treatments by modelling weed abundance per field as a function of both the rotation treatment and the crop or grassland in place, controlling for the plot block (Figure 1B and 1C). As the statistical model includes the effect of the crop in place, the significance of the effect of rotation treatments reported here goes beyond the differences in crop type (i.e., grassland or cereal crop). Crop type also had a strong effect on weed abundance: weed abundance in maize and wheat crops was, on average, about three times higher than that in grassland and barley crops (Figure 1C).

**Figure 1.**
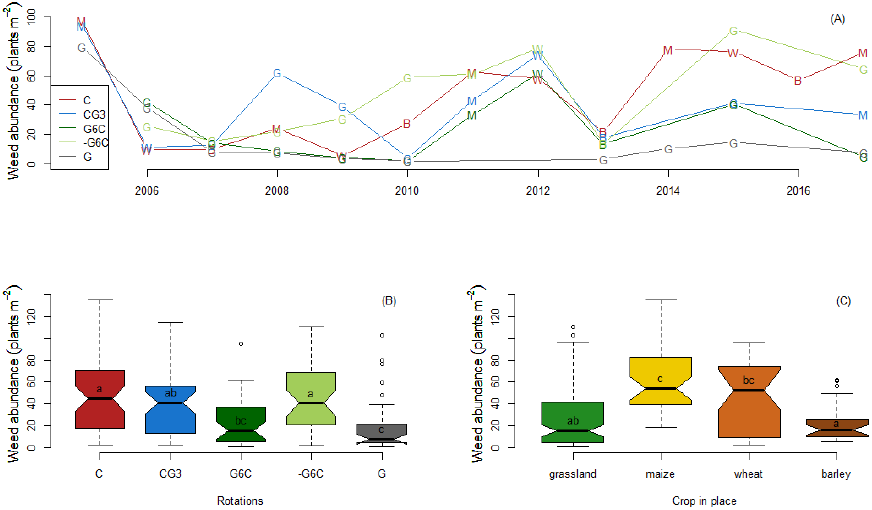
Weed abundance (mean number of plants.m^−2^) dynamics (A), and distribution, by rotation (B) and crop in place (C) during the 12 years of the experiment. The uppercase letters in panel (A) indicate the crop in place: G=grassland, M=maize, W=wheat and B=barley; each point corresponding to the mean abundance over the fields of a rotation. Rotation codes in panel (B): C corresponds to a repeated three-year cereal rotation, G corresponds to continuous grassland and three or six indicates the number of years of well-fertilised grassland and the negative sign (–) corresponds to grassland with a lower level of fertilisation. Different lowercase letters (a, b, c) in panels (B) and (C) indicate significant differences between rotation systems and crops in place in pairwise comparisons, after Holm–Bonferroni adjustment (*P*< 0.05). The boxplots indicate the median (dark line), the 25 and 75% percentiles (limits of the coloured box), and the confidence interval for the median based on an assumption of asymptotic normality of the median (notch). The length of the whiskers corresponds to the minimum of either the distance to the extremes or 1.5 times the length of coloured box, with possible outliers shown as points.

We evaluated the weed-suppressing effect of well-fertilised grasslands in the crops following the temporary grassland, by analysing three years in which the crops were identical in the different rotations. In 2011-2013, all rotations had similar cereal crops in place (except rotation treatment G, the permanent grassland), and all these crops followed a period of grassland phase except for the rotation including only cereal crops. During this three-year period, only rotation treatment G6C, in which the cereal crop followed six years of well-fertilised grassland had a significantly lower weed abundance, 25 to 50% lower than the values obtained for the other rotations (Figure 1A and Table 1). During the next three years (2014 to 2016), grassland was reintroduced in rotation treatments CG3, G6C and -G6C. During this period, weed abundance was lowest in the continuous grassland (G). Well-fertilised grasslands left in place for three or six years (CG3 and G6C) had similar intermediate weed abundances. Finally, weed abundance was higher and similar in the cereal crops (G) and in grassland with a low level of nitrogen fertilisation (-G6C; Table 1).

**Table 1.** Effects of rotations on weed abundance (GLM coefficient “Estimate”) in three year periods of cereal crops after grassland (2011-2013) or grassland after crops (2014-2016), with weed abundance in the cereal crop-only rotation as the reference.

| Rotation | Cereal crop<br>after<br>grassland<br>(2011-2013) | Grassland<br>after cereal<br>crop<br>(2014-2016) |
| --- | --- | --- |

| C* | 0 a | 0 a |
| --- | --- | --- |
| CG3 | -0.141 a | -0.682 b |
| G6C | -0.396 b | -0.724 b |
| -G6C | -0.064 a | 0.124 a |
| G | -1.789 c | -1.866 c |

For rotation codes, C corresponds to the repeated three-year cereal crop rotation, G corresponds to well-fertilised grassland left in place for 3 or 6 years and to grassland with lower levels of fertilisation and \* is the reference rotation. Different lowercase letters (a, b, c) indicate significant differences between rotations in pairwise comparisons with Holm–Bonferroni adjustment ( $P < 0.05$ ).

### 3.2 Comparison of models explaining weed abundance

As rotation treatment had a strong impact on weed abundance over the years, we attempted to model directly the impact on weed abundance of nitrogen fertilisation, grassland duration (i.e., age of grassland) and the crop in place. Strong variability was also observed between years. We therefore also took the major weather variables (i.e., mean precipitation and temperature, and the thermal amplitude of each year) into account. Hypothesising that biomass production might suppress weeds, we compared the explanatory power of the biomass produced in the previous year with that of the aforementioned explanatory factors (AIC differences, Table 2). The biomass of the crop/grassland in the previous year (model 9) largely outperformed all other predictive factors, even combined (model 10).

**Table 2.**
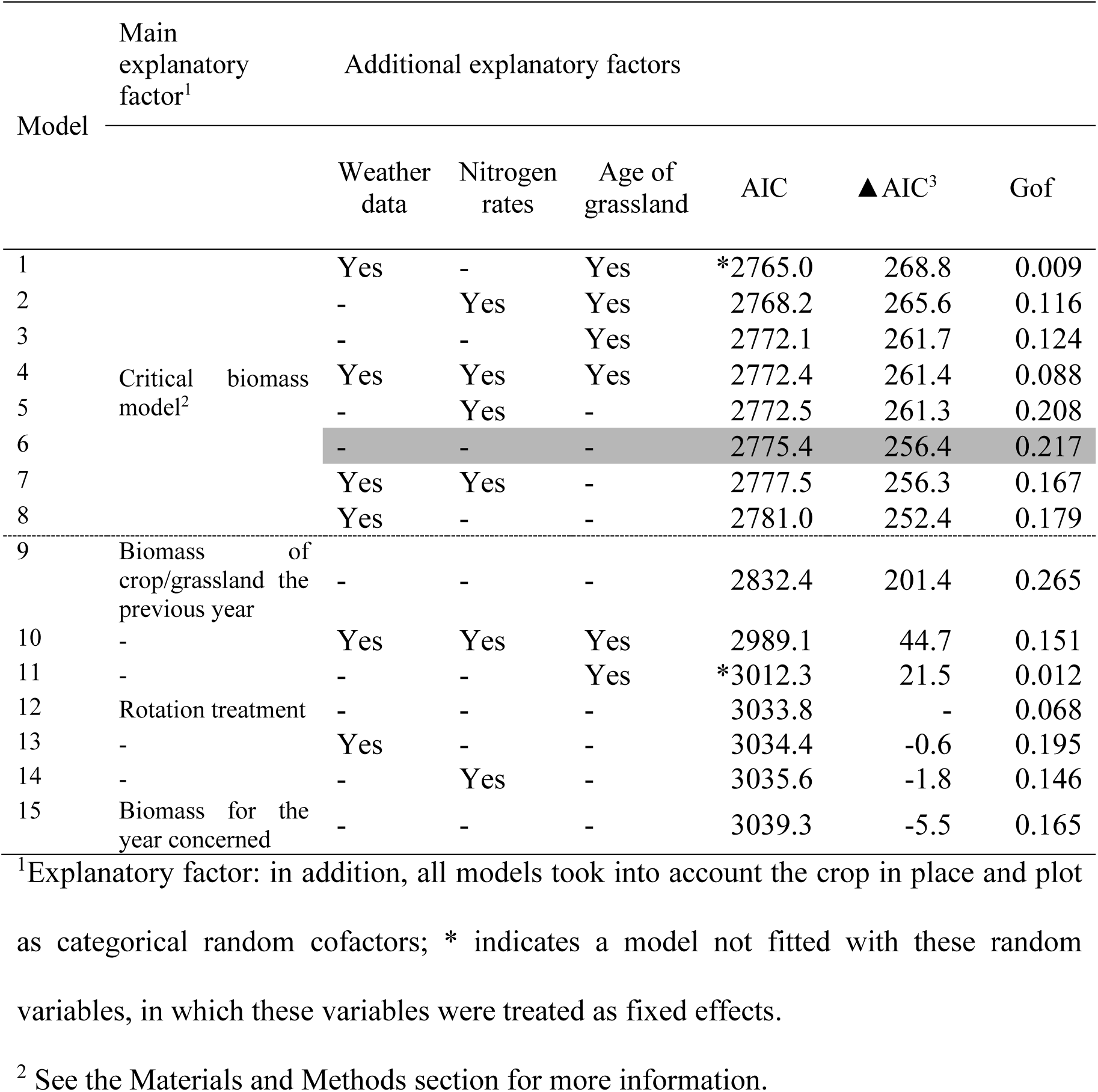
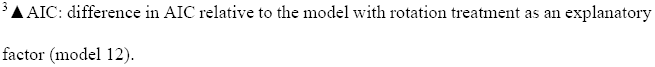
Comparison of the proposed critical biomass model with other models for explaining weed abundance patterns in the systems studied (the proposed model without other variables as explanatory factors is highlighted in grey).

| Model | Main explanatory factor<sup>1</sup> |  | Additional explanatory factors |  |  |  |  |  |
| --- | --- | --- | --- | --- | --- | --- | --- | --- |
|  |  |  | Weather data | Nitrogen rates | Age of grassland | AIC | ▲ AIC<sup>3</sup> | Gof |
| 1 | Critical biomass model<sup>2</sup> |  | Yes | - | Yes | *2765.0 | 268.8 | 0.009 |
| 2 |  |  | - | Yes | Yes | 2768.2 | 265.6 | 0.116 |
| 3 |  |  | - | - | Yes | 2772.1 | 261.7 | 0.124 |
| 4 |  |  | Yes | Yes | Yes | 2772.4 | 261.4 | 0.088 |
| 5 |  |  | - | Yes | - | 2772.5 | 261.3 | 0.208 |
| 6 |  |  | - | - | - | 2775.4 | 256.4 | 0.217 |
| 7 |  |  | Yes | Yes | - | 2777.5 | 256.3 | 0.167 |
| 8 |  |  | Yes | - | - | 2781.0 | 252.4 | 0.179 |
| 9 | Biomass of crop/grassland the previous year |  | - | - | - | 2832.4 | 201.4 | 0.265 |
| 10 | - |  | Yes | Yes | Yes | 2989.1 | 44.7 | 0.151 |
| 11 | - |  | - | - | Yes | *3012.3 | 21.5 | 0.012 |
| 12 | Rotation treatment |  | - | - | - | 3033.8 | - | 0.068 |
| 13 | - |  | Yes | - | - | 3034.4 | -0.6 | 0.195 |
| 14 | - |  | - | Yes | - | 3035.6 | -1.8 | 0.146 |
| 15 | Biomass for the year concerned |  | - | - | - | 3039.3 | -5.5 | 0.165 |

<sup>1</sup>Explanatory factor: in addition, all models took into account the crop in place and plot as categorical random cofactors; \* indicates a model not fitted with these random variables, in which these variables were treated as fixed effects.
<sup>2</sup> See the Materials and Methods section for more information.
<sup>3</sup>▲AIC: difference in AIC relative to the model with rotation treatment as an explanatory factor (model 12).

Finally, we evaluated our proposed critical biomass model and compared it with the other models. The critical biomass model was more than 50 AIC points better than the model with based on the biomass of the crop/grassland in the previous year, and 256.4 AIC points better than the model with rotation treatment as the only explicative factor. We also investigated whether the inclusion of weather data, nitrogen rates and grassland age could improve the critical biomass model. These inclusions improved the model by less than 1%, indicating limited statistical support for the use of these more complex models (≤12.4 AIC points; Table 2) and suggesting that previous crop/grassland biomass production, as described in the critical biomass model, correctly integrates the impact of the other factors on weed abundance.

### 3.3 Evaluation and consistency of the critical biomass model

The weed abundances predicted by our critical biomass model closely matched the observed values (Figure 2A), with a good R^2^ (0.57) and correct ranking (Spearman’s correlation coefficient of 0.75). The bias test revealed a tendency toward overestimation of 4.2% and the mean prediction error (root-mean-square deviation, RMSE) was about 21% (Table 3). No autocorrelation of the residuals over time was detected (Figure 2B), and the residuals displayed no major deviation from a normal distribution (Figure 3C). The statistical estimates can thus be considered accurate.

**Figure 2.**
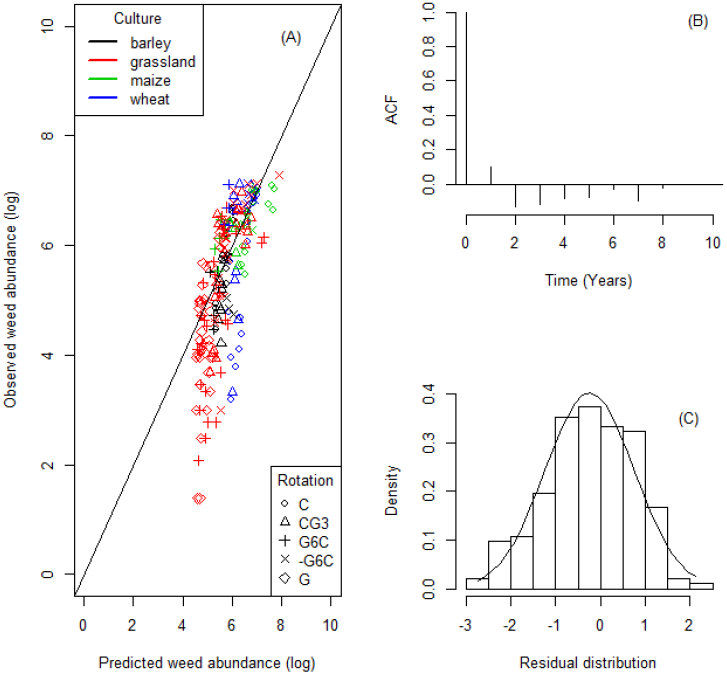
Assessment of the model. Predicted vs. observed weed abundances (log scale) for the crop-grassland rotation treatments (A), autocorrelation (autocorrelation and cross-correlation function, ACF) of the residuals over time (B) and distribution of the residuals of the model (C). Each point represents the weed abundance (over an area of 13 m^2^) per plot and per year on a logarithmic scale. For rotation codes, C corresponds to a repeated three-year cereal rotation, G corresponds to continuous grassland and three or six indicates the number of years of well-fertilised grassland and the negative sign (–) corresponds to grassland with a lower level of fertilisation.

**Table 3.**
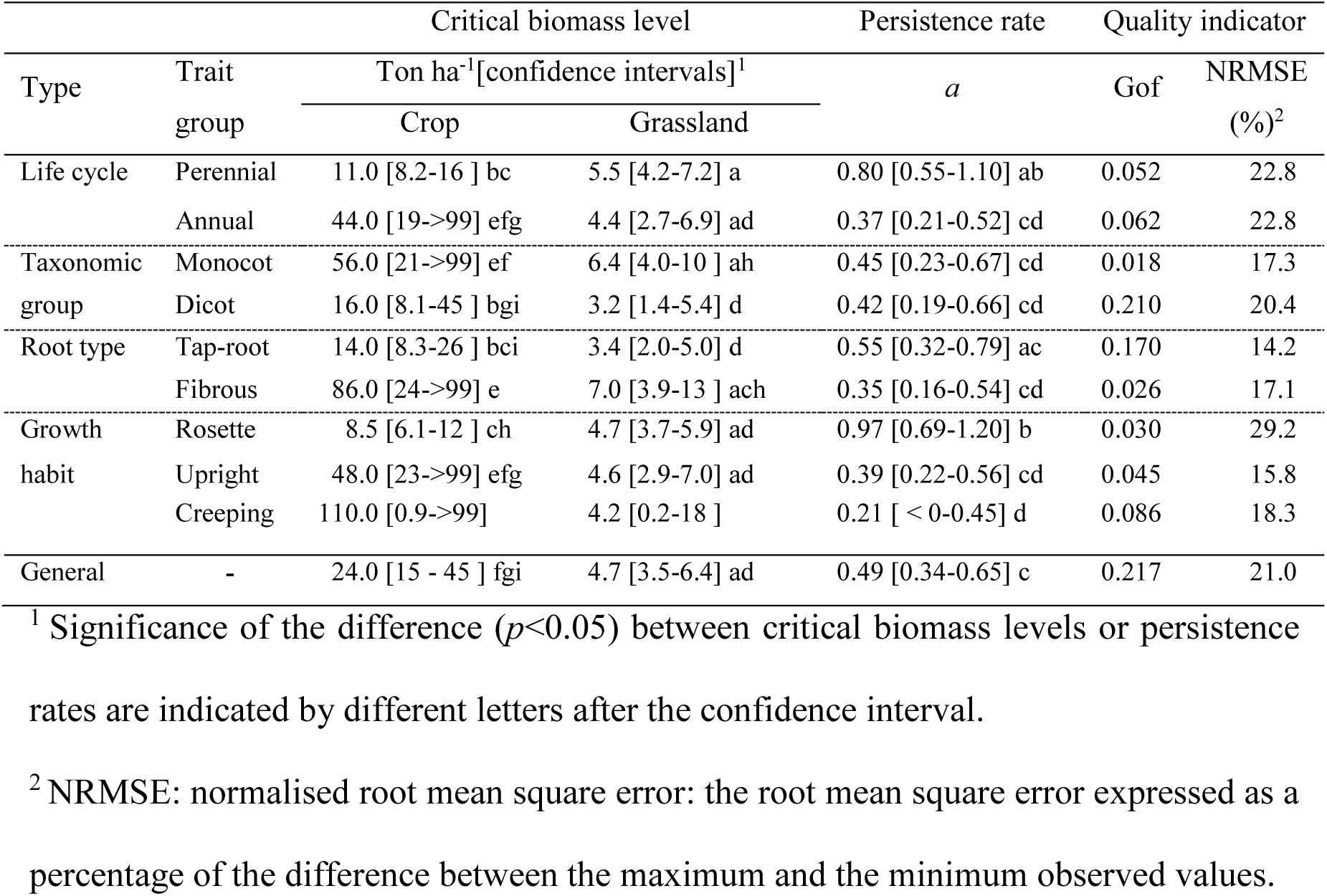
Critical levels of biomass production required to modify (to decrease if higher and to increase if lower) weed abundance the following year, persistence of the effect over time (*a*) and indicators of model quality, as a function of weed trait group.

| Type | Trait group | Critical biomass level |  | Persistence rate | Quality indicator |  |
| --- | --- | --- | --- | --- | --- | --- |
|  |  | Ton ha<sup>-1</sup>[confidence intervals]<sup>1</sup> |  | *a* | Gof | NRMSE (%)<sup>2</sup> |
|  |  | Crop | Grassland |  |  |  |
| Life cycle | Perennial | 11.0 [8.2-16 ] bc | 5.5 [4.2-7.2] a | 0.80 [0.55-1.10] ab | 0.052 | 22.8 |
|  | Annual | 44.0 [19->99] efg | 4.4 [2.7-6.9] ad | 0.37 [0.21-0.52] cd | 0.062 | 22.8 |
| Taxonomic group | Monocot | 56.0 [21->99] ef | 6.4 [4.0-10 ] ah | 0.45 [0.23-0.67] cd | 0.018 | 17.3 |
|  | Dicot | 16.0 [8.1-45 ] bgi | 3.2 [1.4-5.4] d | 0.42 [0.19-0.66] cd | 0.210 | 20.4 |
| Root type | Tap-root | 14.0 [8.3-26 ] bci | 3.4 [2.0-5.0] d | 0.55 [0.32-0.79] ac | 0.170 | 14.2 |
|  | Fibrous | 86.0 [24->99] e | 7.0 [3.9-13 ] ach | 0.35 [0.16-0.54] cd | 0.026 | 17.1 |
| Growth habit | Rosette | 8.5 [6.1-12 ] ch | 4.7 [3.7-5.9] ad | 0.97 [0.69-1.20] b | 0.030 | 29.2 |
|  | Upright | 48.0 [23->99] efg | 4.6 [2.9-7.0] ad | 0.39 [0.22-0.56] cd | 0.045 | 15.8 |
|  | Creeping | 110.0 [0.9->99] | 4.2 [0.2-18 ] | 0.21 [ < 0-0.45] d | 0.086 | 18.3 |
| General | - | 24.0 [15 - 45 ] fgi | 4.7 [3.5-6.4] ad | 0.49 [0.34-0.65] c | 0.217 | 21.0 |

<sup>1</sup> Significance of the difference ( $p<0.05$ ) between critical biomass levels or persistence rates are indicated by different letters after the confidence interval.
<sup>2</sup> NRMSE: normalised root mean square error: the root mean square error expressed as a percentage of the difference between the maximum and the minimum observed values.

According to the critical biomass model of weed abundance, in the studied crop-grassland rotations, the estimated critical levels of biomass corresponding to a stabilisation of weed abundance were 24.3 and 4.7 tonnes dry matter.ha^−1^ for crop and grassland, respectively (Table 3: General). Comparison with actual above-ground biomass over the study — average of 12.1 ton.ha^−1^ for crops (range: 2.7-26.7) and 7.1 ton.ha^−1^ for grassland (range: 0.2-16.6) — highlighted the difficulty reaching the critical biomass level (CBL) for cereal crops, whereas grassland biomass values were generally above the CBL.

### 3.4 Weed trait group responses to previous crop and grassland biomass production

We then investigated whether weeds with different traits were affected differently by the crop and grassland, by fitting the critical biomass model separately to the abundances of different groups of weeds (Table 3). The “critical biomass model” could generally be fitted to the abundances of the different weed trait groups but no convergence of fit could be achieved for weeds with a climbing growth habit.

For all weed traits, the grassland CBL was significantly lower than the cereal crop CBL. The biomass required for weed suppression was two to 10 times higher for cereal crops than for grassland (Table 3). The differences in CBL between weed trait groups were smaller in grassland than in cereal crops. For example, the CBL for cereal crops was almost four times higher for annual weeds than for perennial weeds, whereas the CBL for grassland was 20% lower for annual than for perennial weeds. Consequently, the confidence intervals for grassland critical biomasses also largely overlapped for the different weed trait groups, although some groups had significantly different grassland CBLs. For example, the CBL for monocots in grassland was estimated to be twice that for dicots. The largest difference in crop critical biomass levels was observed for root type, with crop critical biomass levels for tap-rooted weeds only one sixth those for weeds with fibrous roots. By contrast, grassland CBLs for tap-rooted weeds were half those for weeds with fibrous roots, revealing opposite control potentials for grassland and crops for these weed groups. Dicots were more easily outcompeted than monocots by both crops and grassland.

In the critical biomass model, the time required to reduce the impact of a given year under a given threshold is directly proportional to alpha (hereafter called persistence rate). For example, a year with a harvested biomass of 14 ton ha^−1^ and a critical biomass of 10 ton ha^−1^ would result in a large decrease in weed abundance in the following years. This effect would decrease over time and lead to a variation of estimated weed counts of less than 5% after seven years with a persistence rate of 1, whereas it would take three times longer (21 years) to achieve the same result with a persistence rate of 3 (see also alpha_ visualisation_SuppInfo). The estimated persistence rate did not differ significantly between weed trait groups except for perennial life cycle and rosette growth habit, which had persistence rates about twice those of the other weed groups.

## 4. Discussion

In this 12-year field trial, grassland was more competitive against weeds than cereal crops, but this effect was dependent on the grassland receiving sufficient nitrogen fertiliser. Our model suggests that biomass production mediates this conditional suppressive effect and estimates separate critical biomass levels (CBLs) for crops and grassland that must be exceeded for weed suppression in subsequent years. These CBL estimates highlight significant variations of cereal crop and grassland competitiveness according to the traits of the weeds present. The lasting effects of composition are additional to those of the crop in place and are modulated by an estimated persistence rate that differs between weed types.

### 4.1 Factors affecting weed abundance

Weed abundances were lower in grassland than in the maize and wheat phases of the rotations, despite the use of herbicides only during the years in which cereals were grown. This result is consistent with the results of Meiss et al. (2010a), and consistent with grasslands having a greater weed-filtering effect than annual crops. Barley had a low weed abundance similar to that of grassland, probably due to the reported allelopathic activity of barley (Jabran, 2017). In addition to these overall effects of the crop in place on emerged weeds, provided adequate amounts of fertiliser were supplied (i.e., high dry matter production), weed abundance was lower in older grasslands and in subsequent crops, as reported in previous studies (e.g., Schuster 2016; Schuster 2018). We also found that lower levels of nitrogen fertilisation in grassland led to an increase in weed abundance over time. This implies that the mere introduction of grassland for a few years is not sufficient to reduce weed abundance in arable land, and that adequate nitrogen fertilisation is the determinant factor for such a reduction. Nevertheless, it should be borne in mind that the grassland investigated here was a mixture of three grass species (perennial ryegrass, tall fescue and orchard grass) and care must be taken when trying to extrapolate these results to other types of grassland, consisting of monocultures of these species or mixtures of widespread grassland species, grasses or leguminous plants.

### 4.2 Biomass produced as an integrative trait for competitiveness against weeds

The model developed here integrates different rotations and amounts of nitrogen fertilizer into a single meaningful biological variable: biomass production in previous years. Competitiveness, expressed as crop/grassland biomass, may affect different phases of the weed life cycle, including seed germination and emergence, plant survival and vegetative growth, and seed production and survival (Colbach et al., 2014). The abundances reported here were reconstituted from abundance classes susceptible to threshold effects, but these effects should be limited by the use of 13 samples per plot. In any case, the use of more precise data would improve the fit of the model.

One remarkable feature of our model is its handling of multiannual history. The multiplicative impact of previous years on weed abundance is consistent with former observations that the geometric mean growth rate is more appropriate than the arithmetic mean for describing long-term changes in weed abundance in variable growing conditions (Freckleton and Watkinson, 1998). However, this mean is weighted here by the inverse of the number of years elapsed, and persistence rate could be interpreted as the number of years characterising the persistent influence of a given year on the dynamics of the group of weeds considered. This decrease over time in the impact of crop and grassland competition in a given year may depend on the persistence of both the weed seed bank and vegetative organs. Strikingly, perennial weeds had significantly higher estimated persistence rates, consistent with previous observations suggesting that perennial vegetative organs play an important role in weed persistence (Herben et al., 2014). The high persistence rate of rosette weeds was also striking, rosettes weeds are not only perennial (Weed_traits_SuppInfo) but also have, during their vegetative phase, leaves attached just above soil level below cutting height, thus at least partly escaping defoliation at mowing. Nevertheless, additional studies are required to disentangle the components of persistence rate: persistence of the seeds, vegetative organs and asexual reproduction.

### 4.3 Crop and grassland biomass competitiveness and weed traits

Overall, in our crop-grassland rotations, cereal crop biomass did not reduce weed abundance efficiently, with estimated critical levels higher than biomass production in most years. Previous studies reported that crop competition reduced weed growth and fecundity but did not completely prevent weed seed production (Chauhan et al., 2017). As a consequence, weeds tend to thrive in subsequent years in monocrop systems. However, the results of our crop-grassland rotations may be specific to seed rate, row spacing and direction conditions, as previous studies have demonstrated that a modulation of the competitive ability of crops through these factors to achieve effective weed control (Sardana et al., 2017). Further studies are required to determine the impact of such agronomic practices on “critical biomass levels”.

The greater effectiveness of grassland biomass against weeds may be due to the year-round ground cover in grassland and the absence of soil tillage, preserving a living and perennial weed-suppressing mulch (Wiens et al., 2006). Furthermore, a closed canopy is achieved more quickly after mowing in grasslands than during the establishment of cereal crops, resulting in greater resistance to invaders (Milbau et al., 2003).

Fitting the “critical biomass model” separately for each group of weeds as a function of their traits revealed large differences in critical biomass levels between weeds, consistent with previous reports that the traits of weeds are closely related to their competitiveness (Schwartz et al., 2016). Overall, our results are consistent with former observations that grassland suppresses dicot weeds more effectively than monocot weeds, as monocots higher survival rates and grow back more rapidly after grassland cutting (Meiss et al. 2008), an effect possibly reinforced by the use of dicot-specific herbicides in our monocotyledon crops. Our model did not adjust for the climbing growth habit, consistent with a lower susceptibility of weeds with this growth habit to competition, as previously reported (Schuster et al., 2016). Such weeds may grow over the crop and grassland canopy with the help of special structures (e.g. tendrils hooks, twining stems and leaves), enabling them to absorb sunlight with limited competition (Kissmann and Groth, 1997). We also expected perennial plants to be more easily controlled by cereal crops than annual plants, because tillage and weeding could be carried out before the sowing of the crop. Rosette weeds were not very competitive in cereal crops, probably due to their lower levels of access to light relative to upright weeds in these tall, very dense crops. By contrast, critical biomass levels in grasslands were very similar for weeds with these two habits, and the greater competitiveness of rosettes relative to plants with erect growth habits in grassland may also be enhanced by their lower sensitivity to repeated mowing (Meiss et al., 2008).

## 5. Conclusion

This study highlights the importance of crop/grassland competitiveness for weed control. Well-fertilised grasslands are particularly competitive and produce sufficient amounts of biomass to outcompete weeds. The critical biomass model developed here can be used to calculate an intuitive metric of this competitiveness with a simple statistical procedure, paving the way for comparisons of crop and grassland competitiveness against weeds under various pedoclimatic conditions and agronomic practices. It makes use of a limited set of readily available variables, such as crop and grassland rotation schemes and biomass production in previous years. In addition to providing a powerful indicator of crop/grassland competitiveness against weeds, this model could potentially be used to predict weed abundance and to develop environment-friendly weed management strategies.

## Supporting information

analysis R script

## 5. Acknowledgements

This study was partly funded by the Coordenação de Aperfeiçoamento Pessoal de Nível Superior – Brasil (CAPES) – Finance Code 001. It was also supported by the National Research Infrastructure Agro-écosystèmes, Cycles Biogéochimique et Biodiversité. We would like to thank all the technicians of the SOERE-ACBB for their participation in field sampling. We dedicate this publication to Laurent.

## 6. Author contributions

FG, XC and SM set up and oversighted the long term study; SM and XC designed the weed survey and XC, DD, SM and MS collected the data with the SOERE-ACBB technical staff; MS, SM, FG, CB and DD conceived the ideas; CB and MS designed the data analysis methodology, analyzed the data and conceived the model; MS and CB led the writing of the manuscript. FG, SM, AM and XC contributed critically to the drafts. All authors gave final approval for publication.

